# Machine-learning and mechanistic modeling of primary and metastatic breast cancer growth after neoadjuvant targeted therapy

**DOI:** 10.1101/2023.02.22.529613

**Authors:** S. Benzekry, M. Mastri, C. Nicolò, J. ML Ebos

## Abstract

Clinical trials involving systemic neoadjuvant treatments in breast cancer aim to shrink tumors prior to surgery while simultaneously allowing for controlled evaluation of biomarkers, toxicity, and suppression of distant (occult) metastatic disease. Yet such trials are rarely preceded by preclinical testing involving surgery. Here we used a mouse model of spontaneous metastasis after surgical removal to develop a predictive mathematical model of neoadjuvant treatment response to sunitinib, a receptor tyrosine kinase inhibitor (RTKI). Longitudinal data consisted of measurements of presurgical primary tumor size and postsurgical metastatic burden in 128 mice (104 for model training, 24 for validation), following variable neoadjuvant treatment schedules over a 14-day period. A nonlinear mixed-effects modeling approach was used to quantify inter-animal variability. Machine learning algorithms were applied to investigate the significance of several biomarkers at resection as predictors of individual kinetics. Biomarkers included circulating tumor- and immune-based cells (circulating tumor cells and myeloid-derived suppressor cells) as well as immunohistochemical tumor proteins (CD31 and Ki67). Our simulations showed that neoadjuvant RTKI treatment inhibits primary tumor growth but has little efficacy in preventing (micro)-metastatic disease progression after surgery. Surprisingly, machine-learning algorithms demonstrated only limited predictive power of tested biomarkers on the mathematical parameters. These results suggest that presurgical modeling might be an effective tool to screen biomarkers prior to clinical trial testing. Mathematical modeling combined with artificial intelligence techniques represent a novel platform for integrating preclinical surgical metastasis models in outcome prediction of neoadjuvant treatment.

**Major findings:** Using simulations from a mechanistic mathematical model compared with preclinical data from surgical metastasis models, we found anti-tumor effects of neoadjuvant RTKI treatment can differ between the primary tumor and metastases in the perioperative setting. Model simulations with variable drug doses and scheduling of neoadjuvant treatment revealed a contrasting impact on initial primary tumor debulking and metastatic outcomes long after treatment has stopped and tumor surgically removed. Using machine-learning algorithms, we identified the limited power of several circulating cellular and molecular biomarkers in predicting metastatic outcome, uncovering a potential fast-track strategy for assessing future clinical biomarkers by paring patient studies with identical studies in mice.

## Introduction

Neoadjuvant trials in breast cancer (BC) patients involve the administration of systemic treatment for a limited period to treat (and reduce) localized primary tumors prior to surgery. They provide several advantages to assist in novel drug development and translational research (1). For example, neoadjuvant trials can be faster to conduct, require fewer patients, offer the potential for controlled assessment of biological tissue for novel biomarker development, and critically, can potentially limit distant (often occult) metastatic lesions to delay disease recurrence long after treatment has ended (1,2). Yet there are surprisingly few studies that precede neoadjuvant trial design to offer predictive guides to validate drug efficacy, biomarkers, or possible outcomes. In this regard, *in silico* (mathematical) modeling and preclinical *in vivo* testing can be useful. However, mathematical modeling most often occurs as post-hoc analysis in BC trials and studies in mice rarely include clinically relevant systems that capture the complexity of surgical impact on primary/metastatic growth to offer rationalized inclusion of biomarkers in trial design.

To address this gap, here we describe a mathematical modeling framework of neoadjuvant therapy using a combination of preclinical *in vivo* and *in silico* data to provide a predictive platform for treatment outcomes. This extends from our prior work that validated a semi-mechanistic model comparing localized ‘primary’ tumor growth with the growth of spontaneous metastatic disease that occurred after surgery in mouse models of BC (4). We used ‘ortho-surgical’ models (i.e., orthotopic implantation followed by surgical tumor resection) to show that inter-individual variability in the kinetics of metastatic growth could be captured by a (lognormal) distribution of a critical parameter of metastatic aggressiveness, which we termed *‘μ’*. By adding here neoadjuvant treatment to this mathematical modeling framework it allowed us to; 1) formulate and test mechanistic hypothesis about differential effects on primary versus secondary disease, 2) evaluate the impact of biomarkers on metastatic development and 3) investigate the impact of modulating dosing regimen. In addition, machine learning coupled to mechanistic modeling – an approach that we call ‘mechanistic learning’ (5,6) – can screen biomarkers with translational potential and establish predictive models (7). In contrast to classical statistical analysis, machine-learning consists in designing models with predictive power as metric of success, rather than inference properties, and makes use of highly nonlinear models such as regression trees or artificial neural networks (8).

In this study, we examined neoadjuvant treatment with sunitinib, a molecular targeted tyrosine inhibitor (TKI) that can block angiogenesis-associated vascular endothelial growth factor receptors (VEGFRs) along with several other regulators of metastasis (9). Using a VEGFR TKI had several advantages. First, they have a short half-life which allowed us to confine treatment effects to the presurgical period and incorporate multiple variations of treatment dosing, days treated, and time of resection after initial tumor implantation in mice. Second, VEGFR TKIs have shown mixed effects in the perioperative setting in BC (10). While the addition of neoadjuvant sunitinib to chemotherapy improved pathologic complete response rates, long-term results have been more contrasted, with no disease free survival benefit and either none (11) or some (12) overall benefit. As a monotherapy, we and others have demonstrated that robust inhibition of primary tumor growth does not always translate into inhibition of metastasis post-surgically nor improvement in survival (13–15). In our model, we measured multiple cellular and molecular biomarkers at surgery and used machine learning to investigate their predictive power on the mechanistic parameters of our neoadjuvant mathematical metastatic model. Machine learning confirmed a lack of definitive biomarkers, which shows the value of preclinical modeling studies to identify potential failures that should be avoided clinically.

## Materials and Methods

### Cell lines

The human LM2-4^LUC+^ cells are a luciferase-expressing metastatic variant of the MDA-MB-231 breast cancer cell line derived after multiple rounds of *in vivo* lung metastasis selection in mice, as previously described (16). LM2-4^LUC+^ were maintained in DMEM (Corning cellgro; 10-013-CV) supplemented with 5% v/v FBS (Corning cellgro; 35-010-CV), in a humidified incubator at 37oC and 5% CO2. The cell line was authenticated by STR profiling (DDC Medical, USA).

### Drug and doses used

Sunitinib malate (SU11248; Sutent®©, Pfizer) was suspended in a vehicle formulation that contained carboxymethylcellulose sodium (USP, 0.5% w/v), NaCl (USP, 1.8% w/v), Tween-80 (NF, 0.4% w/v), benzyl alcohol (NF, 0.9% w/v), and reverse osmosis deionized water (added to final volume), which was then adjusted to pH 6. The drug was administered at 60 or 120 mg/kg/day orally by gavage as previously described (13,17). The treatment window used in all neoadjuvant studies consisted of a previously optimized 14-day period prior to surgery (13). Within this 14-day period, daily sunitinib (Su) treatment was given either at 60 mg/kg/day (for 3, 7, or 14 days followed by vehicle for 11, 7, or 0 days, respectively), or at 120 mg/kg/day for 3 days followed by 60 mg/kg/day for 0, 4, 8, or 11 days, and vehicle for 11, 7, 3, or 0 days, respectively. An example of an abbreviation in the text includes ‘Su60(14D)’, which means ‘sunitinib at 60mg/kg/day for 14 days. Schematics for all studies are shown in Table S1. Mice treated daily with vehicle for 14 days were used as controls. Detailed analysis and comparisons of these treatment regimens are described in a companion study evaluating treatment breaks on metastatic disease.

### Ortho-surgical model of metastasis

Animal studies were performed in strict accordance with the recommendations in the Guide for Care and Use of Laboratory Animals of the National Institute of Health and according to guidelines of the Institutional Animal Care and Use Committee at Roswell Park Comprehensive Cancer Center (protocol: 1227M, PI: John M.L. Ebos).

*Implantations:* Experimental methodology was extended from previous work using a xenograft animal model of breast cancer spontaneous metastasis that includes orthotopic implantation followed by surgical resection of a primary (termed ‘ortho-surgical’) (4). Briefly, LM2-4^LUC+^(1 × 10^6^ cells in 100μl DMEM) were orthotopically implanted into the right inguinal mammary fat pad (right flank) of 6-8-week old female SCID mice, as described previously (4,13,17). Primary tumor burden was monitored with Vernier calipers using the formula width^2^(length x 0.5) and bioluminescence imaging (BLI) (4,13,17). Neoadjuvant treatments started 14 days before primary tumors were surgically removed at a timepoint (34-38 days post-implantation) previously optimized for maximal distant metastatic seeding but minimal localized invasion (4, 13). The surgeries were planned at specific time points post-implantation to avoid invasion of primary tumor into the skin or peritoneal wall, ensuring that metastatic progression had proceed and minimizing the possibility of surgical cure (4,13). Postsurgical MB was assessed by BLI and overall survival was monitored based on signs of end stage disease as previously described(4,13).
*Exclusion criteria:* Two scenarios represented instances where animals were excluded from treatment studies. First, if complete removal of primary tumor was not surgically feasible because of local invasion or evidence of advanced metastatic spread(4,17). Second, if no primary or metastatic tumor was ever detected by BLI or visual assessment it was assumed there was lack of tumor-take upon implantation (4,13).
*Randomization:* Before treatment initiation animals were randomized by primary tumor size assessed by Vernier calipers to avoid any false results due to unequal tumor burden between groups (18).

### Flow cytometry

Peripheral blood was collected in tubes containing lithium heparin (BD Biosciences; 365965) by orbital bleeding one day before surgical tumor resection. Non-specific binding was blocked with normal mouse IgG (Invitrogen; 10400C) incubated with whole blood, followed by incubation with an antibody mix. After staining, cells were fixed in a lyse/fix solution (BD Biosciences; 558049), while red blood cells were lysed. Samples were analyzed with a LSR II low cytometer (Becton Dickinson), while data were acquired with FACSDiva software (Becton Dickinson) and analyzed with FCS Express 6 (DeNovo software).

### Circulating tumor cells (CTC)

The antibody mix for CTC detection of human CTCs in animal models contained a rat anti-mouse CD45 (30-F11) antibody conjugated to PE (Biolegend; 103106) and mouse anti-human HLA conjugated to AlexaFluor 647 (Biolegend; 311416). CD45 staining with a rat anti-mouse CD45 conjugated to FITC (Invitrogen; MCD4501) was used to eliminate any mouse blood cells, whereas human HLA was used to identify CTC (human LM2-4^LUC+^). For a positive control, LM2-4^LUC+^ cells were trypsinized, washed with PBS, and stained for both CD45 and HLA. LM2-4^LUC+^ were used to define the CTC gate.

### Circulating myeloid-derived suppressor cells (MDSC)

The antibody mix for detection of MDSCs contained a rat anti-mouse CD45 (30-F11) antibody conjugated to PE (Biolegend; 103106), a rat anti-mouse Ly-6G/Ly-6C (Gr1) (RB6-8C5) antibody conjugated to PE-Cy7 (BD Pharmingen; 552985), and an rat anti-mouse CD11b (M1/70) antibody conjugated to eFluor450 (eBioscience; 48-0112). Mouse CD45 staining was used to select only leukocytes, and CD11b and Gr1 were used to define the granulocytic and monocytic MDSC.

### Immunofluorescence

Resected tumors were frozen on dry ice in cryo-embedding compound (Ted Pella, Inc; 27300), sectioned, and fixed in a 3:1 mixture of acetone:ethanol. Non-specific binding was blocked with 2% BSA in PBS, followed by staining with antibody mix containing rabbit anti-mouse Ki67 antibody (Cell Signaling Technologies; 12202) and rat anti-mouse CD31 antibody (Dianova; DIA-310). Detection of primary antibodies was achieved using FITC conjugated goat anti-rabbit IgG (BD Pharmingen; 554020) and Cy3 conjugated goat anti-rat IgG (Invitrogen; A10522). Samples were counterstained with DAPI (Vector; H-1500) and mounted with a hard-set mounting medium for fluorescence. Random images from each section were obtained with a Zeiss AxioImager A2 epifluorescence microscope at 200x magnification, and analyzed with ImageJ. CD31+ cells (% area) and Ki67+ cells (% cells) were quantified automatically using macro functions, whereas Ki76+/CD31 + cells (proliferating endothelial cells) were quantified manually.

### Mechanistic model of ortho-surgical metastasis

For untreated animals, we previously validated a mechanistic model for description of pre-surgical primary tumor and post-surgical metastasis kinetics in the ortho-surgical LM2-4^LUC+^ animal model (4). Briefly, metastatic development is decomposed into two main processes: growth and dissemination.

Growth of the primary and metastases follow the Gomp-Exp model (19):

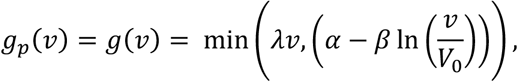

where *g_p_* and *g* denote the growth rates of the primary and secondary tumors, respectively. Parameter *λ* limits the Gompertz growth rate to avoid unrealistically fast kinetics for small sizes and is given by the *in vitro* proliferation rate, assessed previously (4). Parameters *α* and *β* are the Gompertz parameters, and *V*_0_ is the size of one cell (in units of mm^3^ for the PT, and photons/seconds for the metastases). The PT volume, *V_p_*(*t*) thus solves

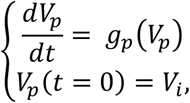

with *V_i_* the volume corresponding to the number of cells injected (= 1 mm^3^ based on the conversion rule 1 mm^3^ ≃ 10^6^ cells (20)).

Dissemination occurs at the following volume-dependent rate (4):

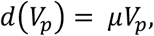

where parameter *μ* can be interpreted as the daily probability that a cell from the PT successfully establishes a metastasis (4).

The metastatic process was described through a function *ρ*(*t,v*) representing the distribution of metastatic tumors with size *v* at time *t*. It solves the following initial boundary value problem (21):

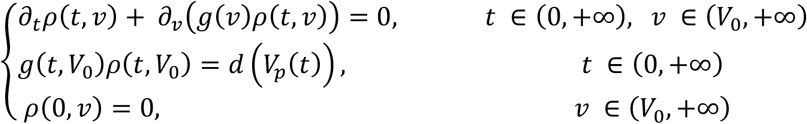

The first equation derives from a balance equation on the number of metastases; the second equation is a boundary condition for the rate of newly created metastases; the third equation is the initial condition (no metastases exist at the initial time).

The total MB at time *t* was then given by

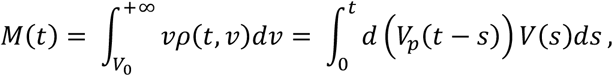

which can be solved efficiently through the use of a fast Fourier transform algorithm (22). In the previous equation, *V*(*s*) represents the volume reached by a metastatic tumor after a period of time s from its emission, when growing with growth rate *g*.

### Mechanistic model of neoadjuvant targeted therapy

Using this model, we next incorporated the effects of systemic therapy. This new model includes neoadjuvant sunitinib treatment and assumes that the drug reduces the primary tumor growth rate by a term proportional to its concentration, *C*(*t*) (Norton-Simon hypothesis (23)):

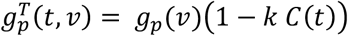

where *k* is a parameter of drug efficacy. As no pharmacokinetic data was available, we used a kinetics-pharmacodynamics (K-PD) approach. Namely, we considered that the drug concentration decays exponentially after each dose,

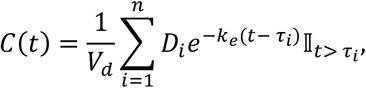

where *D_i_* indicates the dose administered at time *τ_i_*. The volume of distribution *V_d_* and the elimination rate constant *k_e_* were fixed to the values reported in (24). Inclusion of treatment effect on metastatic growth was considered in the model development phase; however, this led to model predictions which could not explain the behavior of the experimental data. Therefore, the final model considered that sunitinib did not affect growth of metastases during neoadjuvant treatment.

### Calibration of the mathematical metastatic model and parameter estimation

Following previously established methodology (4), the mathematical metastatic model was fitted to the experimental data using a nonlinear-mixed effects modeling approach (25). Briefly, this consists in modeling inter-animal variability by assuming a parametric distribution for the model parameters. All individual PT and MB longitudinal data could then be pooled together in a population model, whose parameters were estimated by likelihood maximization (25). In mathematical terms, if 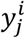 denotes the observation (primary tumor size or metastatic burden) in animal *i* at time 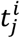 and 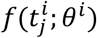 denotes the model value in an animal with parameter set 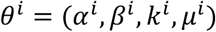, the statistical model linking the model to the observations writes:

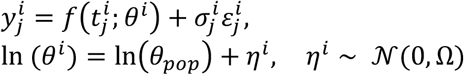

where 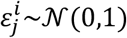 is a gaussian noise for measurement error. The parameters *θ_pop_* and Ω characterize the entire population. The observed data were log-transformed and a proportional error model was used, that is

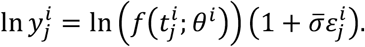

For the vector of individual parameters, a log-normal distribution with full covariance matrix was assumed. Maximum likelihood estimates of the population parameters were obtained using the Stochastic Approximation of Expectation-Maximization (SAEM) algorithm implemented in the nlmefitsa Matlab function (26). PT and MB data were fitted simultaneously for vehicle and sunitinib-treated animals. Visual predictive checks (VPC), individual fits and standard diagnostic graphical tools based on individual parameters were used for evaluating the adequacy of the different model components.

### Machine learning algorithms

Effects of covariates on the model parameters were assessed using linear regression and a number of machine learning regression techniques (partial least squares, artificial neural networks, support vector machines and random forest models) using the R caret package (27,28). Except for the random forest models, data were centered and scaled prior to modeling. Tuning parameter values of the regression models were selected to minimize the root mean squared error (RMSE) using five replicates of a 10-fold cross-validation. If *θ^i^* are the true values and 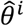 the predicted ones, the RMSE is defined by:

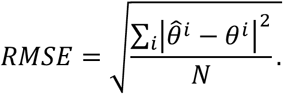

## Results

### *In vivo/in silico* modeling of neoadjuvant treatment

We have previously developed mathematical parameters of untreated spontaneous systemic metastatic breast cancer using orthotopic tumor implantation and surgical resection (i.e., ‘ortho-surgical’) models (4). Using individual longitudinal presurgical primary tumor (PT) and postsurgical metastatic burden (MB; tracked by bioluminescence) data, we previously established that the metastatic potential parameter *μ* quantifies inter-individual variability after surgical resection of the PT (4). In addition, our results validated the use of the mathematical model to simulate pre- and post-surgical metastatic development. Here, using a xenograft model with highly-metastatic human breast cells expressing luciferase (LM2-4^LUC+^), we evaluated PT and MB data from 128 mice that received multiple doses and durations of neoadjuvant sunitinib treatment over a 14 days period (schematic shown in Fig 1A; see Table S1 and methods for treatment details).

**Figure 1:**
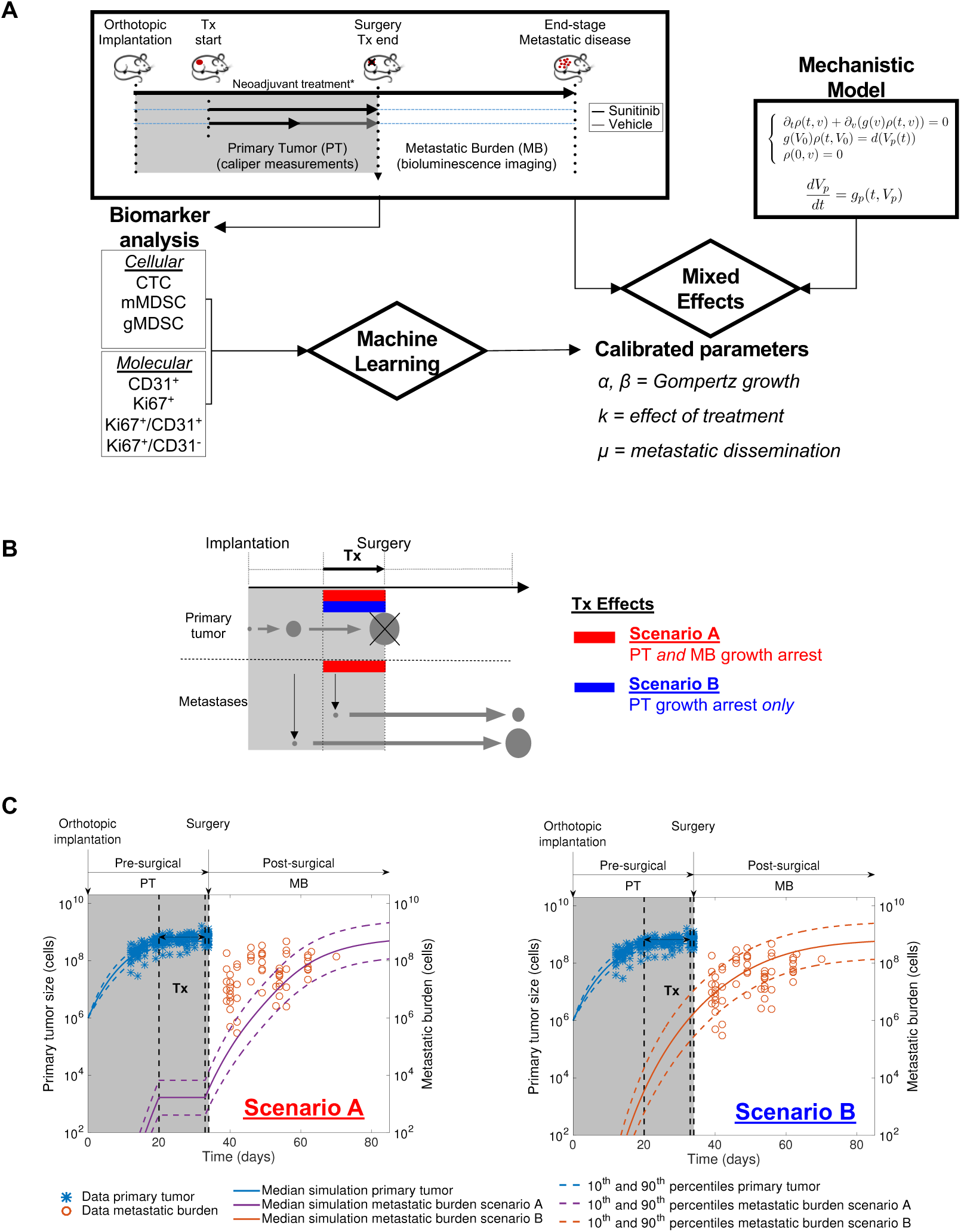
Mathematical modeling reveals differential effects of neoadjuvant sunitinib treatment on primary tumor and metastatic growth. **(A)** Schematic of the study. Data from an ortho-surgical, human xenograft animal model of neoadjuvant sunitinib breast cancer treatment were fitted using a mixed-effects statistical framework. This provided calibrated parameters for each animal. Machine learning algorithms were used to assess the predictive power of molecular and cellular biomarkers to predict the metastatic dissemination parameter μ and quantify metastatic aggressiveness. Biological and numerical parameters quantified at end of therapy and at time of surgery were implemented into a survival model. **(B)** Schematic of tested hypotheses of the effect of neoadjuvant sunitinib Tx on primary tumor and metastatic growth and dissemination through mechanistic mathematical modeling. Scenario A = growth arrest on both primary and secondary tumors. Scenario B = growth arrest on primary tumor only. **(C)** Predicted simulations of Scenarios A and B using parameters calibrated from a previous study [Benzekry et al., Cancer Res, 2016] involving untreated (vehicle) animals only. Data plotted here (LM2-4^LUC+^ bioluminescent human breast cancer cells orthotopically injected in mice) was not used to estimate the model parameters. *Tx, treatment; PT, primary tumor; MB, metastatic burden. *See methods for additional details on animal experiments, treatment dose and duration, and mechanistic model*.

### Simulations of neoadjuvant sunitinib targeted treatment therapy (NATT) suggests limited effect on metastasis growth

We have previously utilized ortho-surgical metastasis models to evaluate the impact of multiple VEGFR TKIs on PT and MB progression (13,14). These studies uncovered that neoadjuvant targeted therapy (NATT) yielded differential effects with suppression of presurgical PT growth not consistently translating into reduction in postsurgical MB nor improvement in survival (13). This effect could result from two phenomena that are mixed in MB quantification: 1) metastatic growth suppression and 2) reduction of metastatic spread as a consequence of primary tumor size reduction. To disentangle the two and qualitatively assess the effect of NATT, we generated predictive model scenarios under two assumptions. In ‘scenario A’, NATT would have growth-arresting effects on both PT and MB, while in ‘scenario B’ NATT would have effect only on PT (schematic shown in Fig 1B). To test this, we used our previously calibrated ortho-surgical model of pre- and postsurgical growth using LM2-4^LUC+^ tumor cells grown in SCID mice (4), and only set either both growth rates *g_p_* and *g* (Scenario ‘A’) or *g_p_* only (Scenario ‘B’) to zero during NATT. Scenario ‘A’ clearly failed to describe the data (Figs 1C and S1), whereas Scenario ‘B’ interestingly demonstrated good accuracy given that simulations were pure predictions that did not make use of the data for parameter estimation (Figs 1C and S1). These results demonstrate a differential effect of NATT on growth of primary and secondary tumors and suggest that a mathematical model of NATT in our breast cancer ortho-surgical animal model should not include anti-growth effect on metastasis.

### Calibration and validation of a kinetics-pharmacodynamics (K-PD) model for pre- and post-surgical disease after neoadjuvant sunitinib therapy

To further link dose and scheduling to response, we developed a K-PD metastatic model of NATT using a defined neoadjuvant treatment window (14 days) containing multiple treatment periods (3, 7, 11, 14 days), doses (60mg and 120mg), and time of surgery after tumor implantation (Day 34 or 38) (see Table S1 and methods for details). Following our findings above, we only adapted the PT growth rate *g_p_* from (4), using the Norton-Simon hypothesis for PT anti-growth effect of NATT (23). Estimates of the model parameters are reported in Table 1 and demonstrate high practical identifiability (relative standard error ≤ 17%), likely owing to the large number of subjects in the population fit. Confirming our previous results (4), the metastatic potential parameter *μ* was found to vary significantly amongst individuals (largest coefficient of variation). Visual predictive checks for both vehicle group and treated groups demonstrated accurate goodness-of-fit both at the population (Figs 2A and S2) and individual (Figs 2B and S3) levels. In addition, model predictions in independent data sets not used for parameter calibration, with distinct time of surgery (day 38 versus day 34) and drug regimens, were in good agreement with the data (Fig S4). Further model diagnostic plots demonstrated no clear misspecification of the structural and residual error model (Fig S5). Distributions of the empirical Bayes estimates were in agreement with the theoretical distributions defined in the statistical model (Fig S6). Moreover, the *η*-shrinkage was less than 20% for each parameter, meaning that the individual parameter estimates and the diagnostic tools based on them can be considered reliable (29). Finally, correlations found between the estimated random effects (Fig S7) confirmed the appropriateness of a full covariance matrix in the assumed distribution of the individual parameters. Together, these results show the validity of our mathematical model to simulate PT and MB kinetics under a wide range of NATT administration regimens, which can thus be employed to explore *in silico* the quantitative impact of possible NATT schedules.

**Table 1:**
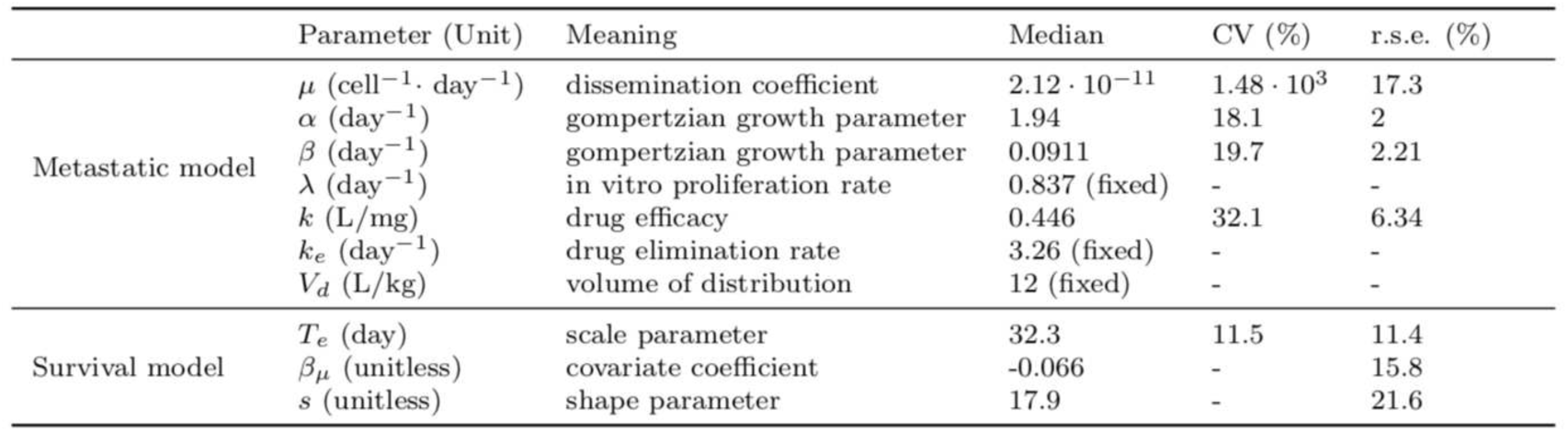
Parameter estimates of the metastatic and survival models obtained by likelihood maximization via the SAEM algorithm. In the survival model, log(mu) has been included as covariate on the scale parameter Te: *log(Te) = log(Te,pop)+ β log(μ) + η*. Abbreviations: CV, coefficient of variation computed as the ratio of the standard deviation and the median of the estimated parameter distribution; r.s.e., residual standard error.

|  | Parameter (Unit) | Meaning | Median | CV (%) | r.s.e. (%) |
| --- | --- | --- | --- | --- | --- |
| Metastatic model | $\mu$ ($\text{cell}^{-1} \cdot \text{day}^{-1}$) | dissemination coefficient | $2.12 \cdot 10^{-11}$ | $1.48 \cdot 10^3$ | 17.3 |
| | $\alpha$ ($\text{day}^{-1}$) | gompertzian growth parameter | 1.94 | 18.1 | 2 |
| | $\beta$ ($\text{day}^{-1}$) | gompertzian growth parameter | 0.0911 | 19.7 | 2.21 |
| | $\lambda$ ($\text{day}^{-1}$) | in vitro proliferation rate | 0.837 (fixed) | - | - |
| | $k$ (L/mg) | drug efficacy | 0.446 | 32.1 | 6.34 |
| | $k_e$ ($\text{day}^{-1}$) | drug elimination rate | 3.26 (fixed) | - | - |
| | $V_d$ (L/kg) | volume of distribution | 12 (fixed) | - | - |
| Survival model | $T_e$ (day) | scale parameter | 32.3 | 11.5 | 11.4 |
| | $\beta_\mu$ (unitless) | covariate coefficient | -0.066 | - | 15.8 |
| | $s$ (unitless) | shape parameter | 17.9 | - | 21.6 |

**Figure 2:**
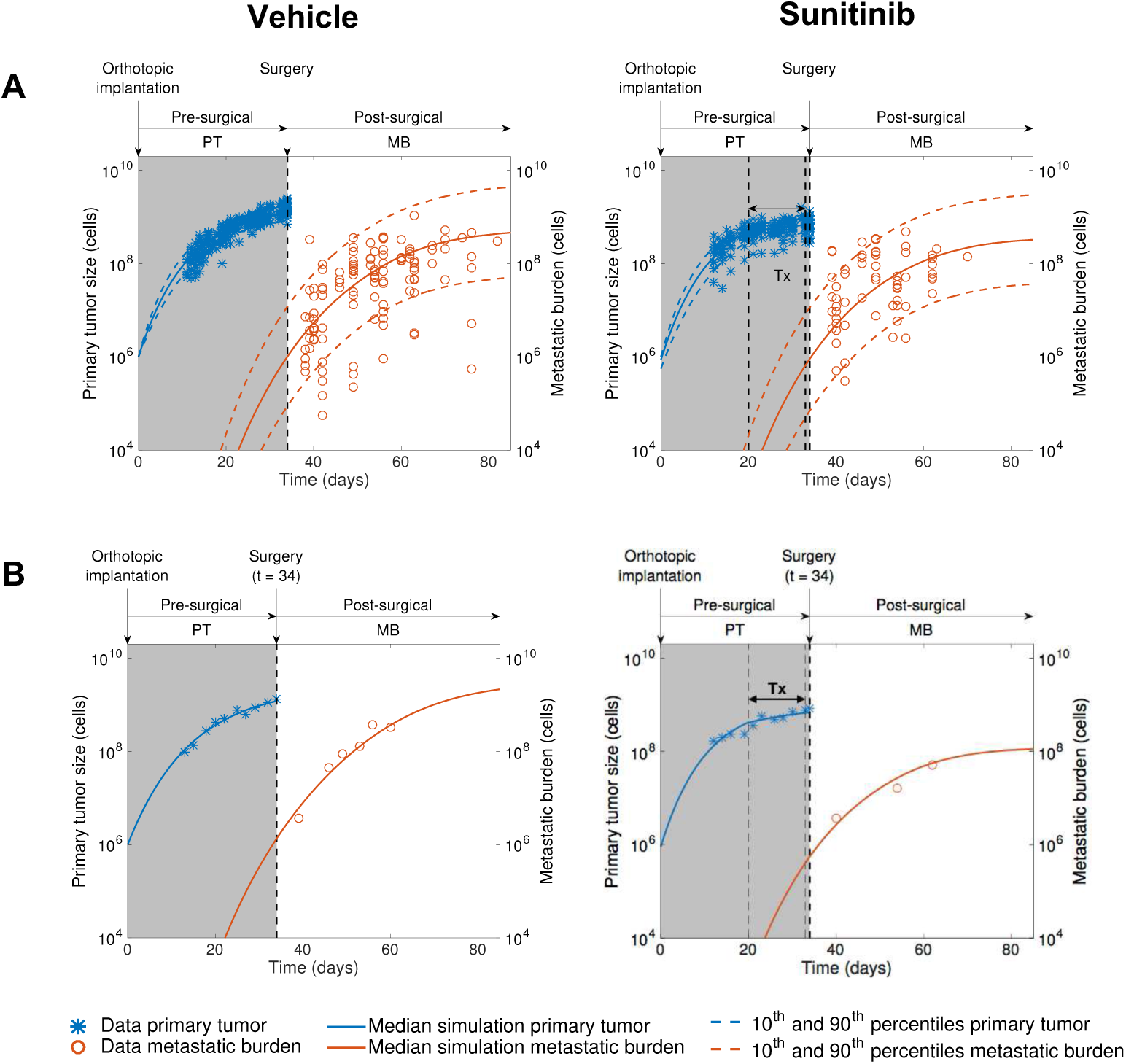
Calibration and validation of a kinetics-pharmacodynamics (K-PD) mathematical model for neoadjuvant sunitinib treatment effect on pre- and post-surgical tumor growth. Pre- and postsurgical growth of LM2-4^LUC+^ human metastatic breast carcinomas were measured in multiple groups involving different neoadjuvant treatment modalities (doses and durations). The mathematical model was fitted to the experimental data using a mixed-effects population approach (n=104 animals in total). **(A)** Comparison of the simulated model population distribution (visual predictive check) for vehicle and neoadjuvant sunitinib treatment (60mg/kg/day) 14 days prior to surgery. **(B)** Examples of individual dynamics. *Tx, treatment; PT, primary tumor; MB, metastatic burden*.

### Simulations of NATT duration reveal contrasted impact on PT size reduction and metastasis-free survival

The overall impact of NATT is the combination of i): PT debulking (which in turn reduces metastatic spread from the PT), and ii) an increased risk of metastatic relapse due to delayed removal of the PT. To quantify the impact of NATT duration on these two opposite aspects, we ran simulations of our calibrated model for 0 to 18 days NATT and three dose levels (60, 120 and 240 mg/kg, see Fig 3). First using only the typical population estimates of the parameters (median individual), we found an important increase in postsurgical MB for long NATT: final values ranged from 2.73 × 10^8^ to 7.52 × 10^8^ cells for NATT durations from 0 to 18 days (176% increase), respectively, at the 60 mg/kg dose level (Fig 3A). This is consistent with our model where NATT does not affect metastatic growth, thus delaying surgery can only increase MB. This was less important in higher dose levels (125% and 48.5% increases for 120 mg/kg and 240 mg/kg, respectively). To study the impact of inter-individual variability, we leveraged our mixed-effects framework to perform population simulations and quantify final outcome. Namely, we simulated 1000 virtual individuals and recorded the percent changes in PT size at the end of NATT. Fig 3B shows the resulting median PT percent changes, together with an area covering 80% of the population. In addition, we calculated a risk of metastatic relapse from the resulting simulation of MB kinetics. To do so, we considered as MB relapse threshold the 30^th^ percentile of the control population MB at 85 days (considered to be an approximation of long term), to mimic the human situation in which 30% of breast cancer patients with localized disease undergo metastatic relapse (30). This threshold then allowed us to compute the percent of subjects having metastatic relapse in the virtual populations, under varying NATT duration (Fig 3B, circled line). For 60 mg/kg and 120 mg/kg doses, metastatic relapse risk is predicted to increase drastically when delaying PT removal too long. However, for a 240 mg/kg dose (or for virtual subjects with increased sensitivity to treatment), increase in metastatic relapse risk is more moderate, since a prolonged NATT is associated with large decrease of the PT size, thus reducing the source of metastasis. Together, these results illustrate how our mathematical model, informed from preclinical data of an NATT ortho-surgical model, can provide informative quantitative simulations of the impact of treatment schedules. Our findings suggest a moderate to detrimental impact of long sunitinib NATT at low dose, in breast cancer.

**Figure 3:**
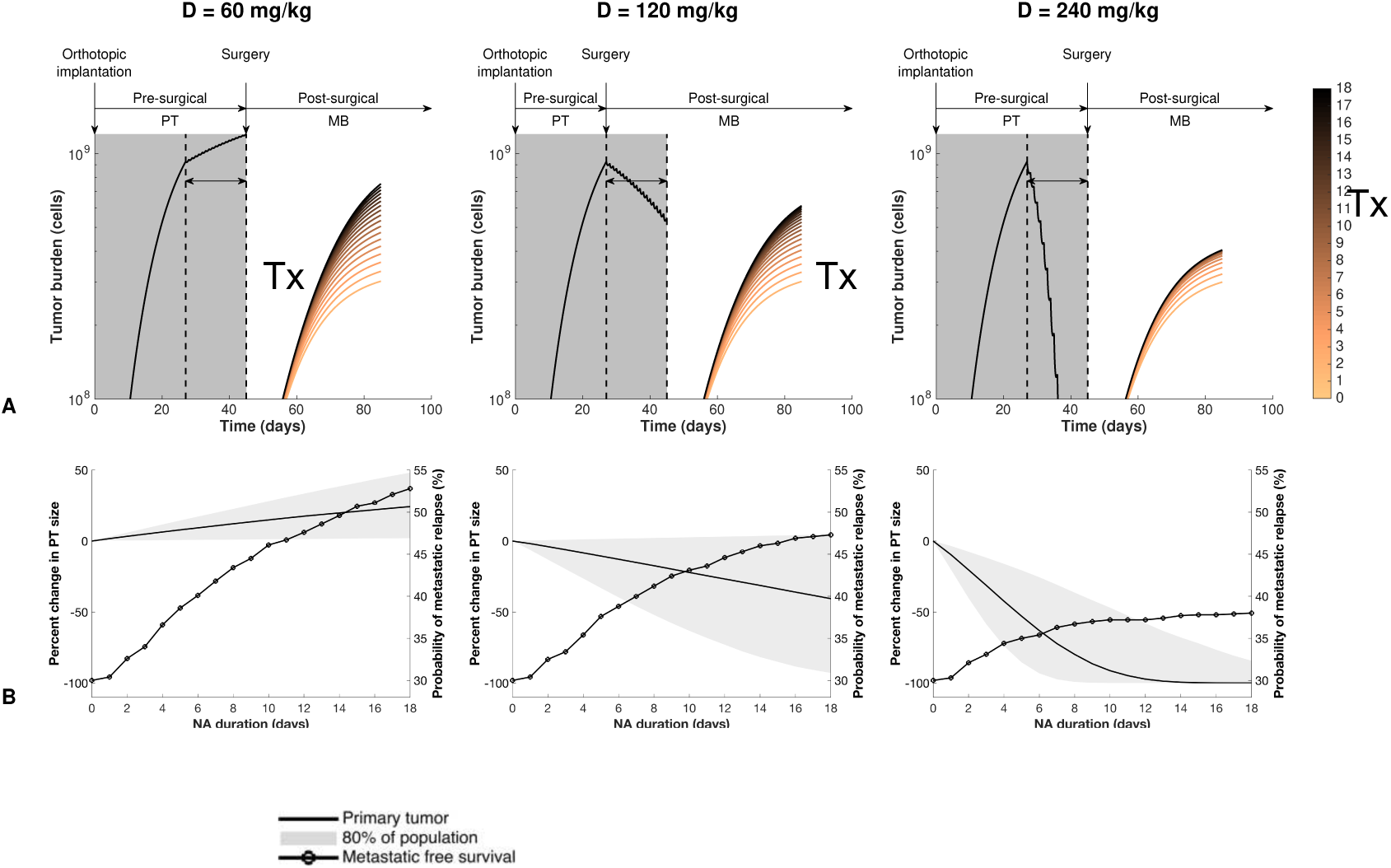
Simulations of varying neoadjuvant treatment duration quantify contrasted impact on primary tumor size reduction and risk of metastatic relapse. Using model parameters calibrated from data of our ortho-surgical animal model of breast cancer neoadjuvant treatment (NAT), simulations were conducted for a duration of NAT varying between 0 (light color) and 18 (dark color) days, for three dose levels (60 mg/kg, 120 mg/kg and 240 mg/kg). **(A)** Predicted simulations of pre-surgical primary tumor and post-surgical metastatic kinetics. Primary tumor growth curves are not distinguishable because they are all superimposed until time of surgery. **(B)** Population-level predictions of final primary tumor size (solid line and grey area) and probability of metastatic relapse as functions of duration of neoadjuvant treatment, which delays surgical removal of primary tumor (circled line). Inter-individual variability simulated from population distribution of the parameters learned from the data (n = 1000 virtual subjects).

### Machine learning for prediction of the metastatic aggressiveness parameter *μ* from biomarkers at surgery

Next, we wanted to determine whether biological parameters at the time of PT surgery but after NAT had stopped could be utilized as predictive biomarkers of postsurgical MB after treatment cessation. These biomarkers included immunohistochemical molecular protein measurements of resected PT for cell proliferation (Ki67) and blood vessel (CD31) markers in resected PTs (Fig 4A; example shown), blood-based cellular measurements of circulating myeloid derived stromal cells (MDSCs) (Fig 3B), and circulating tumor cells (CTCs) from 66 animals (Fig 4B and 4C, respectively; examples of flowcytometric gating shown). We investigated whether these molecular and cellular biomarkers may parallel the observed variability in the mathematical parameters, in particular *μ*, whose large variability indicated potential animal subpopulations of variable metastatic potential values. We first examined correlations between biomarkers in order to identify potential redundancies in the data (Fig 4B). High correlations were found between Ki67 and Ki67+/CD31- (r = 0.979, p < 10^−12^) and CTC and gMDSC (r = 0.678, p = 3.95 · 10^−10^). Next, we investigated the value of these measurements as predictive biomarkers of the mechanistic parameters: *α* and *β* capture growth kinetics, *k* the effect of treatment and *μ* metastatic dissemination. Fig 4B shows correlations between biomarkers and the parameter estimates. As the individual growth parameters *α* and *β* were highly correlated (r = 0.997, p < 10^−5^), we used the Gompertz tumor doubling time at the volume *V_i_* = 1 mm^3^ to assess the impact of covariates on the tumor growth parameters. It is defined by 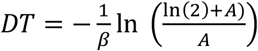, with 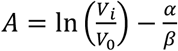. A weak correlation was found between log (*DT*) and mMDSC levels (Fig 4E, r = 0.275, p = 0.0257). However, none of the available biomarkers was found to correlate either with *μ* or log (*μ*) (Fig S8). Next, partial least squares and a number of different machine learning regression algorithms were tested in order to identify possible relationships between covariates and individual estimates of the metastatic potential parameter (shown in Fig 1A schematic). These included neural networks, support vector machines and random forest models (31). Cross-validation results for the RMSE of the final regression models were compared against the intercept-only model (the constant model were predictions are the same for all animals, given by the median value in the population, *μ_pop_*). As shown in Figs 4F and 4G, none of the fitted models had RMSE or R^2^ significantly different from the intercept-only model. Lowest RMSE was achieved by the intercept-only model. Values of R^2^ ranged from 0.133 to 0.199 across the models, with the highest value reached by the conditional random forest model. Prediction error on ln(*μ*) ranged from 9.83% ± 10.7% for the best model (conditional random forests, mean ± std) to 10.6% ± 11.3% for the worse (random forests), which was not superior to predictive power of the intercept-only model (9.71% ± 10.1%). Plotting the observed versus predicted values (Figs 5 and S9) confirmed that the fitted algorithms were unable to explain the variability of parameter *μ*. Together, these results demonstrate that the biomarkers considered in this study have limited predictive power for metastatic potential as defined by *μ*.

**Figure 4:**
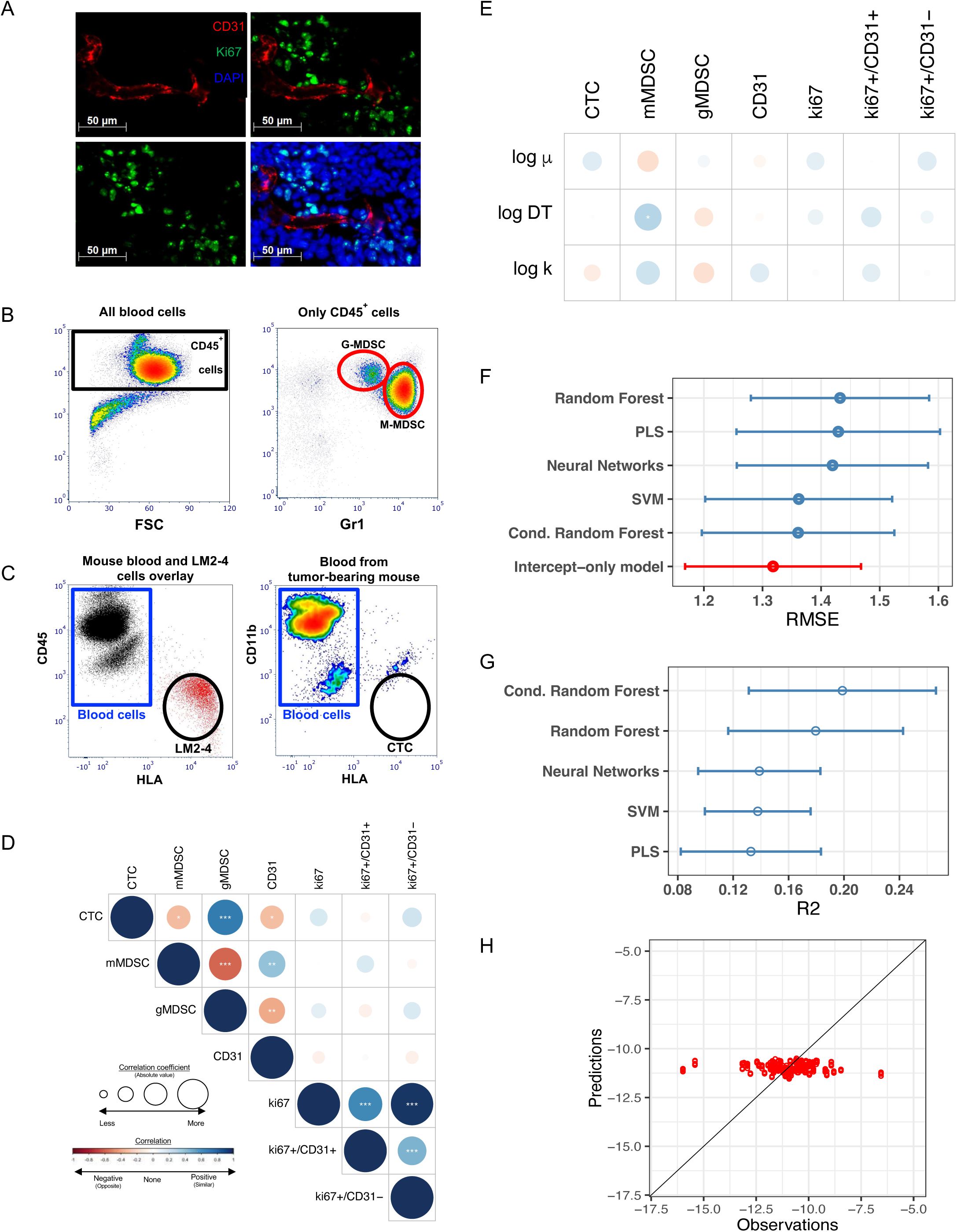
Use of machine learning algorithms based on presurgical molecular and cellular markers to predict metastatic dissemination parameter ‘μ’. **(A-C)** Examples of molecular and cellular biomarker analysis. **(A)** Proliferating endothelial cell identification by immunofluorescence. Tissue sections from resected tumors were stained with antibodies against mouse CD31 (red) and mouse Ki67 (green) and counterstained with DAPI (blue). Single channel and merged images are shown. Yellow arrows show proliferating endothelial cells which were counted manually. **(B)** Myeloid-Derived Suppressor Cells (MDSC) quantification by flow cytometry. Whole blood was stained with anti-mouse antibodies for CD45, CD11b, and Gr1. After selection of CD45 positive cells MDSCs were analyzed based on CD11b and Gr1 levels. Monocytic-MDSC (M-MDSC) are CD11b+/Gr1high and granulocytic-MDSC (G-MDSC) are CD1 1 b+/Gr1 Medium. Examples of MDSC in untreated and treated animals are shown. (**C)** CTC quantification by flow cytometry. CTCs for xenografts were identified using anti-human HLA. Blood was stained with anti-mouse CD45 and anti-human HLA. Blood and LM2-4 cell samples were overlaid in a dot plot to identify and create the gates for CTCs. Once the gates were created CTC were identified in blood of tumor-bearing mice. **(D)** Pearson correlation coefficients between biomarkers. Blue (resp. red) color indicates positive (resp. negative) correlation, with size of the circle proportional to the R^2^ correlation coefficient. *\*p<0.05, **p<0.01, ***p<0.001*. **(E)** Univariate correlations between the biomarkers and the mathematical parameters. DT = doubling time. **(F)** Cross-validated Root Mean Square Error (RMSE) across different machine learning regression models (see methods) utilizing the values of the biomarkers for predicting log(*μ*). To assess the significance of the covariate in the models, RMSE were compared against the value of this metric obtained using a only-intercept model. Bars are 95% confidence intervals. Shown in red is the model with lowest RMSE. PLS = Partial Least Squares. SVM = Support Vector Machines **(G)** Cross validated R2 with 95% confidence intervals. **(H)** Predictions versus observations for the conditional random forest algorithm.

## Discussion

A large part of *in vivo* studies in experimental therapeutics focus on the effect of treatments on isolated tumors and few make use of metastatic animal models (32). However, we and others have previously shown that differential effects occur on the primary tumor and the metastases for some anti-cancer drugs, such as the multitargeted tyrosine kinase inhibitor sunitinib (13,14,33). Similarly, apart from efforts focusing on evolutionary dynamics of metastasis that do not make use of longitudinal data on size kinetics (34), few quantitative mathematical models exist for metastatic development (4,21,35,36), and none has been quantitatively validated for systemic therapy beyond theoretical considerations (35,37,38). In previous work we first established such a mathematical model featuring natural metastatic development and surgery of the primary tumor, but no systemic treatment (4). This was a critical step before being able to model the effect of systemic treatments such as NATT where treatments are limited and long-term benefits are presumed but difficult to quantify as disease recurrence can happen years after surgery, or not recur at all. In the current study, we extended our mathematical model to examine NATT with the RTKI sunitinib by using longitudinal data of 128 mice (more than four times more than previous studies (4,36)). Such large number of subjects and tightly controlled experimental conditions (genetically identical animal background, cell origin, treatment periods, etc..), resulted in precise estimates of the model parameters. Together our results represent an idealized system for predicting treatment impact and novel biomarker identification that could assist in trial design prior to testing in patients.

Our results using sunitinib showing efficacy in reducing primary tumor growth but not metastasis mirror our previous report with two ortho-surgical animal models where we found that NATT with sunitinib and axitinib (another VEGFR TKI) did not always limit metastatic disease after surgery, despite clear antitumor effects on localized disease (13). This represents a challenge observed clinically with RTKIs where, despite decades of potent tumor reducing effects in mouse models, efficacy in patients with metastatic disease could be underwhelming. Using our mathematical model to simulate distinct biological scenarios, we demonstrated that the effect of the drug on tumor growth could differ between primary and secondary sites. Conversely, model simulation predictions (with no fitting involved) of a scenario where metastatic development during NATT was only altered by primary tumor size shrinkage was in excellent agreement with the data.

These findings could be explained by the fact that the primary tumor (in the mammary fat pad) and the secondary tumors (mostly in the lungs) would rely on different growth mechanisms, especially at small sizes. Supporting this explanation, a study showed that metastasis relied more on vessel co-option rather than angiogenesis, thus providing them a mechanism of resistance to VEGFR TKIs therapy (39). Beyond NATT, our model predicts limited efficacy of sunitinib in the postsurgical setting, because metastases would likely be similarly small and rely on similar growth mechanisms. Interestingly, recent experimental results in mice confirmed this prediction where, using a similar metastatic experimental system of triple negative breast cancer, adjuvant sunitinib did not improve survival (40). The mechanistic model of NATT validated here provides a valuable tool to explore the impact of the treatment schedules on response and relapse. Simulating varying durations and doses of NATT, we found that long durations of NATT could significantly increase the risk of metastatic relapse when PT response was moderate. Further, our model provides the computational basis to analyze the impact of various NATT dosing regimen in terms of sequence, breaks and frequency, which is the topic of a companion work.

For breast cancer patients diagnosed with localized disease, predicting the risk and timing of distant metastatic relapse is a major clinical concern (41–43). Accurate ways to predict the extent of invisible metastatic disease at diagnosis and risk of future metastatic relapse could help to personalize perioperative therapy protocols, and avoid highly toxic therapies to patients with low risk of relapse (42). However, only two risk models (44,45) have met the AJCC criteria for prognostic tool quality so far (46), and both rely on classical Cox regression survival models. Recently, we have developed a mechanistic approach to metastatic relapse prediction (47). However, this work did not include the impact of NATT not any systemic treatment. The mathematical model that we validated here on animal data combined with the methodology developed in (47) lays the groundwork for applications in the clinical NATT setting. It could further refine individual predictions of metastatic relapse in breast cancer by providing surrogate markers of long-term outcome additional to pathologic complete response (3). Indeed, the NATT time period represents an invaluable window of opportunity to gather both longitudinal data (such as kinetics of tumor size or pharmacodynamic marker, or circulating DNA from liquid biopsies) and one-time biomarkers from tumor tissue (2). Here, we propose that mathematical models could form the basis of digital tools able to integrate this multi-parametric and dynamic data into predictive algorithms of both long-term outcome and disease sensitivity to systemic therapy in case of distant relapse.

In the era of artificial intelligence (48), it is to be expected that an increasing number of such prognosis models will appear, combining advances in cancer biology (e.g. molecular gene signatures (42,49)) and imaging (50,51) with algorithmic engineering. Recent years have witnessed the generalization of methods going beyond classical statistical analysis, grouped by the generic term of machine learning (ML) (52). However, these techniques have not been applied to preclinical data from targeted therapy. Here, we proposed an approach to combine ML with mechanistic modeling that consists of using biomarkers at surgery to predict individual mathematical parameters and subsequently postsurgical metastatic evolution. Multiple cellular and molecular biomarkers were measured at the time of surgery, either by immunohistochemistry or flow cytometry. These constituted candidate features for ML prediction of the critical parameter *μ*, which we found as being the major driver of inter-subject metastatic variability. We found overall that these biomarkers contained only limited predictive power of *μ*, suggesting that alternative biomarkers should be explored in future preclinical and clinical studies. This contrasts with reports showing Ki67 as significantly associated with risk of metastatic relapse (53). It might be due to the fact that Ki67 is a proliferation marker (54), which should rather be predictive of *α* or the doubling time. In fact, such correlation was observed between Ki67+/CD31+ and DT (Fig 3E), as well as clinical work using our modeling approach (47). Paired with early clinical trials, our *in vivo/in silico* approach could have translational value to inform the screening of biomarkers.

Important limitations of our study are that we only analyzed data from one tumor type (triple negative breast cancer), one cell line in one, immune-depressed, animal system and one drug. On the other hand, this is a necessary prerequisite to control as much as possible the heterogeneity in the data, which still remains substantial despite a tightly controlled experimental setting. Such conditions ensure robust test of biological assumptions underlying our mathematical models and, eventually, refutation of unplausible ones (here, that primary and secondary growth would be equally suppressed by NATT). Nevertheless, to address these limitations, we conducted a companion study in similar ortho-surgical kidney cancer systems, with two cell lines (SN12-PM6-N and RENCA, respectively of human and murine origin), immune-competent animals (Balb/c mice for the RENCA cells), and two VEGFR TKIs (Sunitinb and Axitinib) used in the clinical setting to treat kidney cancer patients. In addition, we investigated in depth the impact of breaks and high-dose “bursts” during NATT. The mathematical model developed on the basis of the one in this study allowed to i) demonstrate and quantify post-NATT PT growth rebound and ii) quantify the impact of such dosing regimen variations on post-surgival metastatic development.

Given the increasingly diverse arsenal of systemic anti-cancer therapies available with the approval of immune-checkpoint inhibitors, optimal treatment sequence (5,55–57) and dosing regimen (58,59) are becoming crucial issues. Our model could be used and extended to guide the rational design of treatment schedules and modes of combination of immunotherapy with another systemic drug, before preclinical or clinical testing. For immunotherapy, the model would need to be developed further and at least include an additional systemic variable representing the immune system. Immuno-monitoring quantifications could provide an invaluable source of longitudinal data to feed mechanistic models (60). In addition, response to neoadjuvant therapy could be used to predict which patients are more likely to benefit from adjuvant therapy (12). Combining artificial intelligence techniques with mechanistic modeling, our modeling methodology offers a way to perform such predictions quantitatively and possibly personalize therapeutic intervention.

## Supplementary Table

**Table S1:** Neoadjuvant treatment schedules and doeses. Animal groups showing treatment schedules and dosing during a presurgical neoadvjuant period of 14 days.

| Group | N | Duration (Days) |  |  | Surgery (DPI) | Modeling | Abbrev. |
| --- | --- | --- | --- | --- | --- | --- | --- |
|  |  | Sunitinib (mg/kg/day) |  |  |  |  |  |
|  |  | 120 | 60 | 0 |  |  |  |
| 1 | 6 | 0 | 0 | 14 | 38 | Training | Veh. |
| 2 | 21 | 0 | 0 | 14 | 34 | Training | Vech. |
| 3 | 21 | 0 | 14 | 0 | 34 | Training | Su60(14D) |
| 4 | 6 | 0 | 7 | 7 | 34 | Training | Su60(7D) |
| 5 | 15 | 0 | 3 | 11 | 34 | Training | Su60(3D) |
| 6 | 15 | 3 | 11 | 0 | 34 | Training | Su120(3D)/Su60(11D) |
| 7 | 20 | 3 | 0 | 11 | 34 | Training | Su120(3D) |
| 8 | 6 | 3 | 11 | 0 | 38 | Validation | Su120(3D)/Su60(11D) |
| 9 | 6 | 3 | 8 | 3 | 38 | Validation | Su120(3D)/Su60(8D) |
| 10 | 6 | 3 | 4 | 7 | 38 | Validation | Su120(3D)/Su60(4D) |
| 11 | 6 | 3 | 0 | 11 | 38 | Validation | Su120(3D) |

## Supplementary Figures

**Figure S1.**
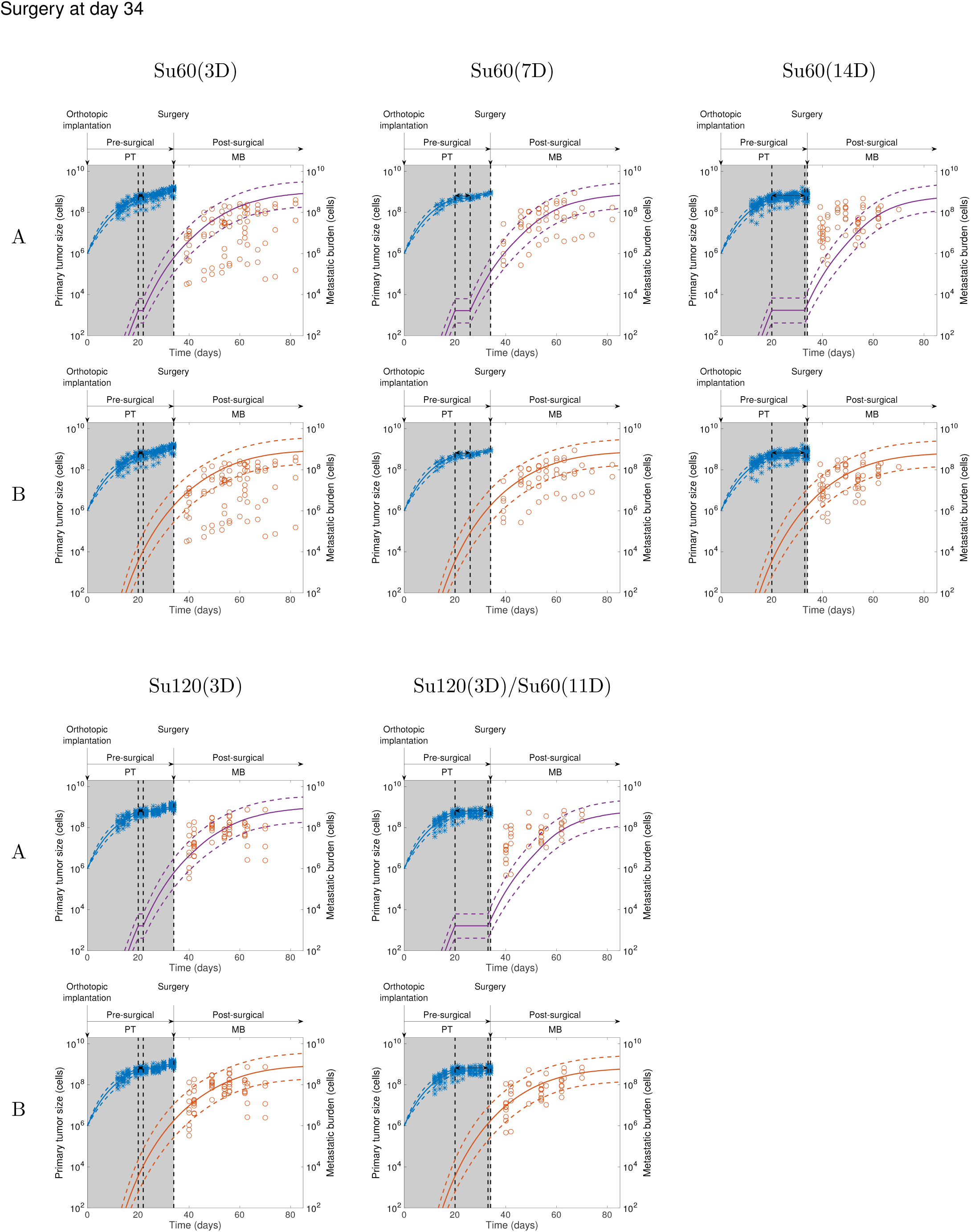
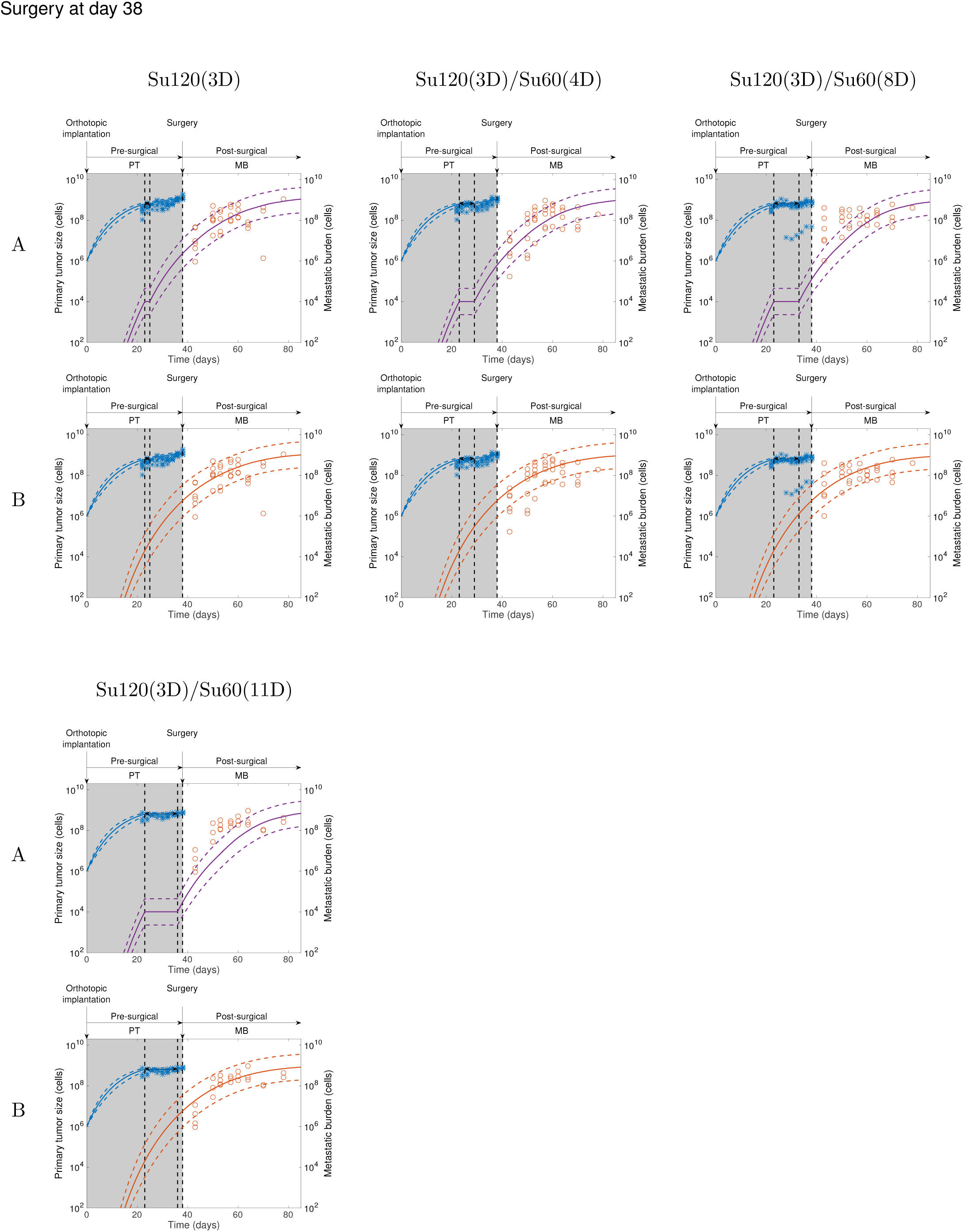
Comparison of simulation of therapy (A) vs no therapy (B) on metastases.

**Figure S2.**
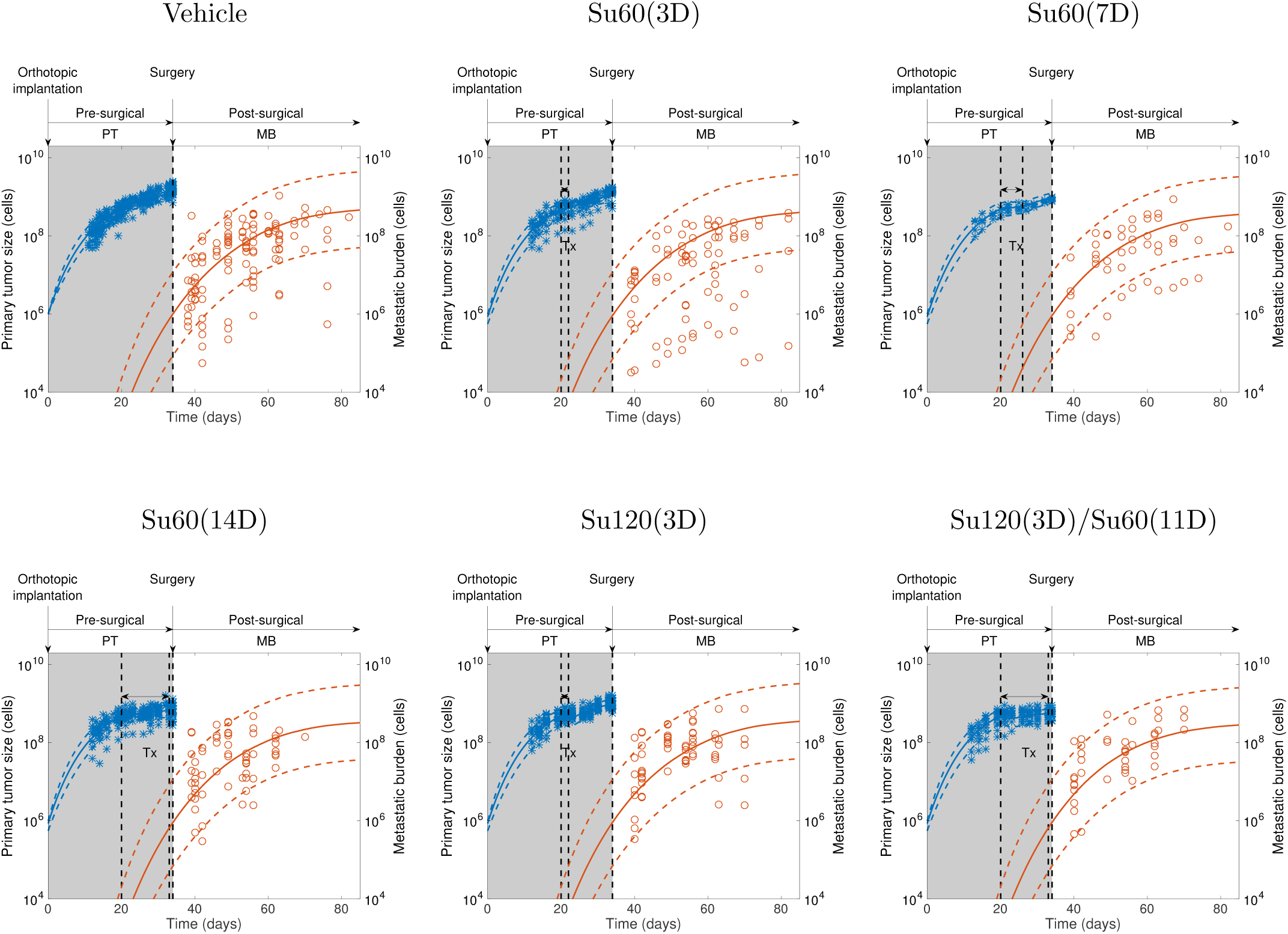
Population fits of all the groups used to calibrate the model parameters (surgery at day 34)

**Figure S3.**
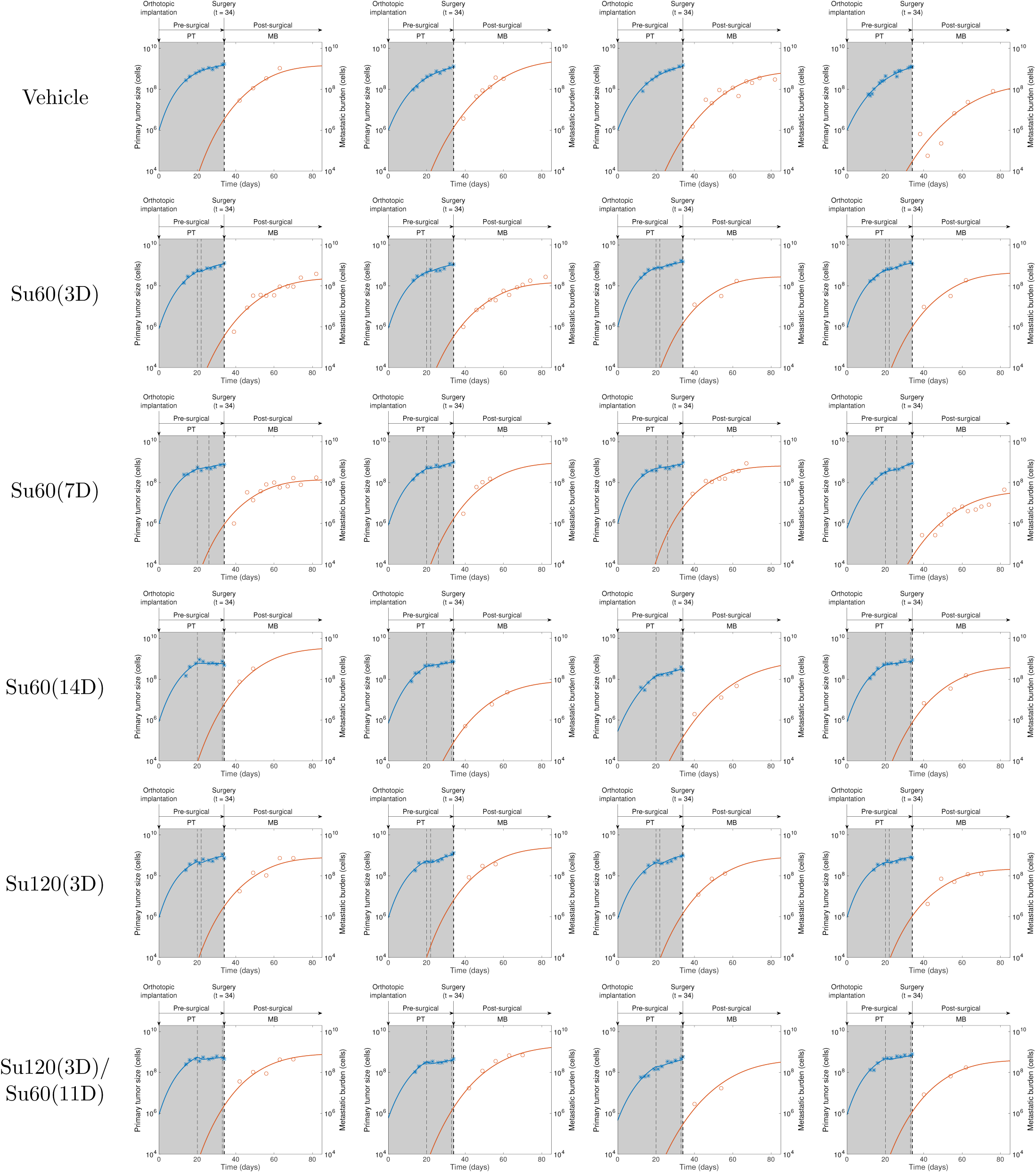
Representative individual fits of the model for Sunitinib-treated animals.

**Figure S4.**
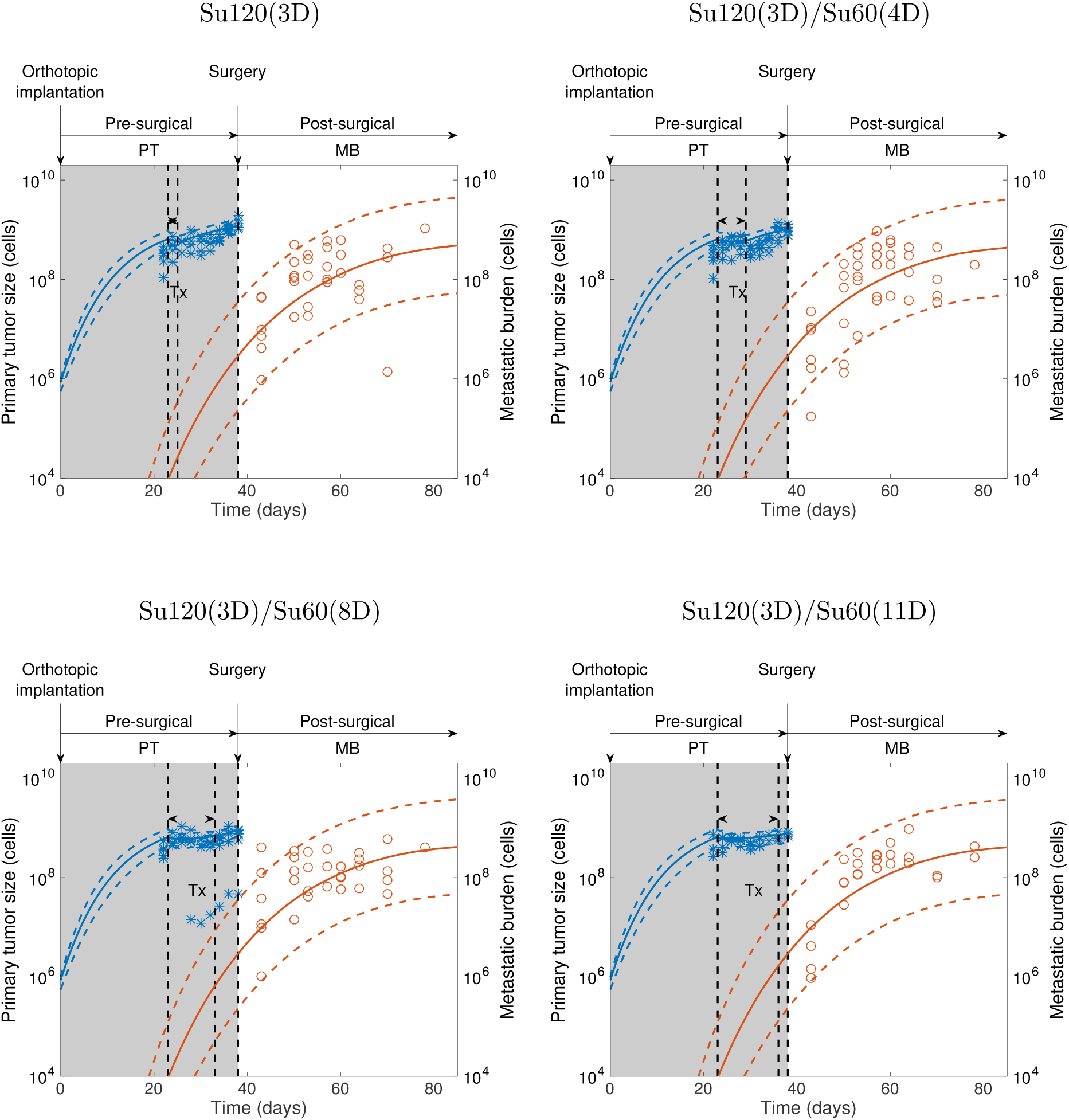
Model predictions in independent datasets (surgery at day 38)

**Figure S5.**
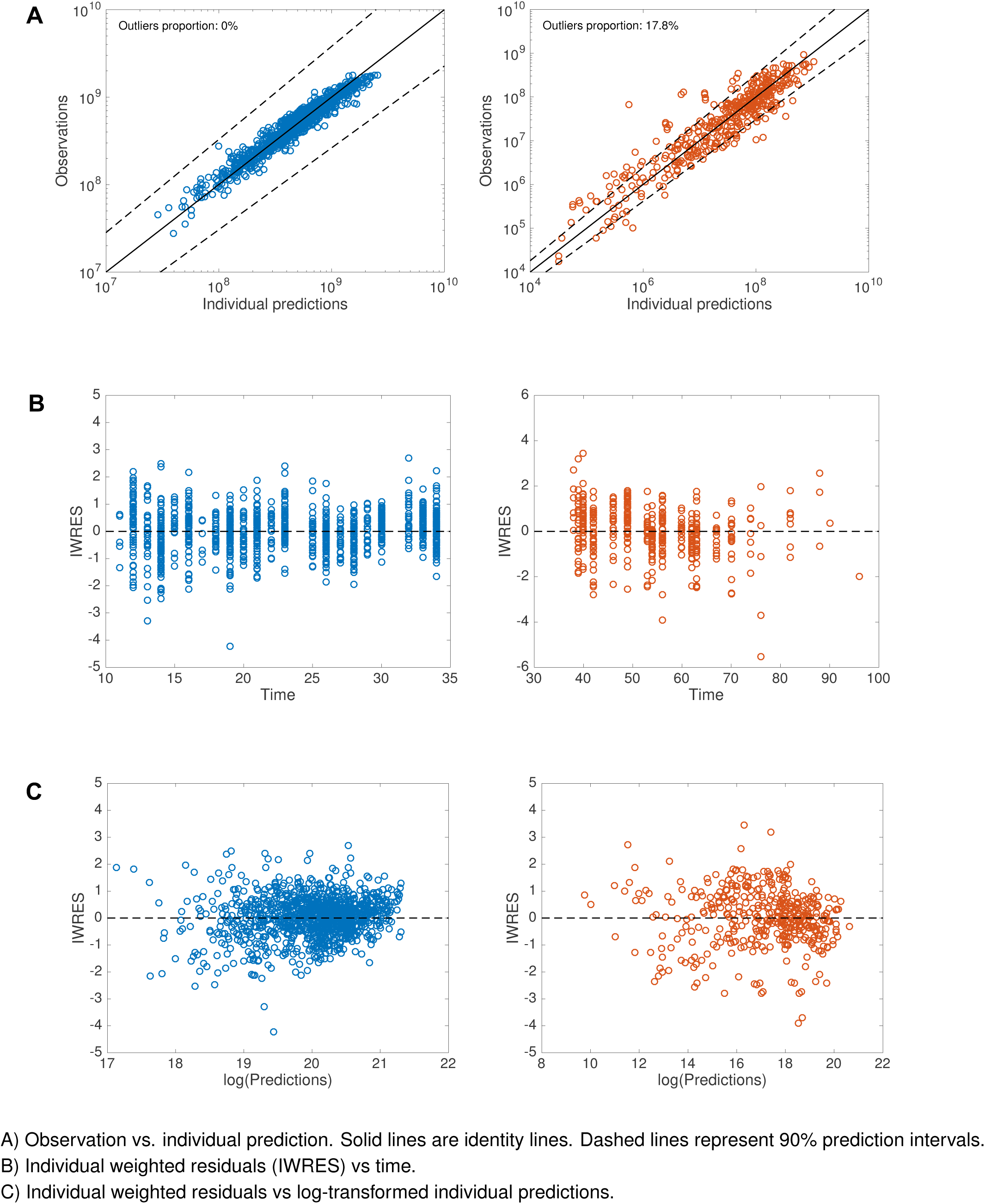
Model diagnostic plots. A) Observation vs. individual prediction. Solid lines are identity lines. Dashed lines represent 90% prediction intervals. B) Individual weighted residuals (IWRES) vs time. C) Individual weighted residuals vs log-transformed individual predictions.

**Figure S6.**
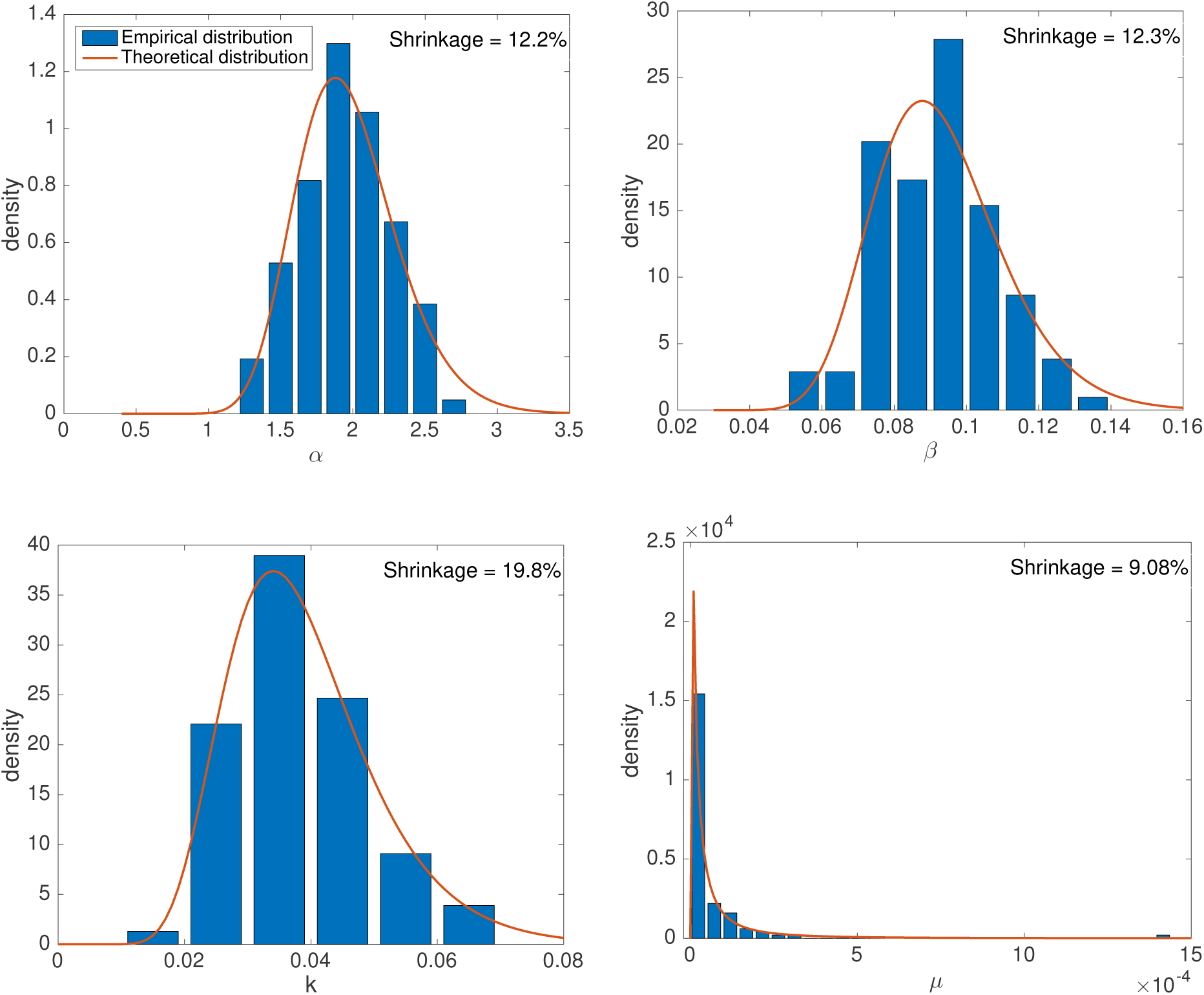
Distribution of the individual parameters.

**Figure S7.**
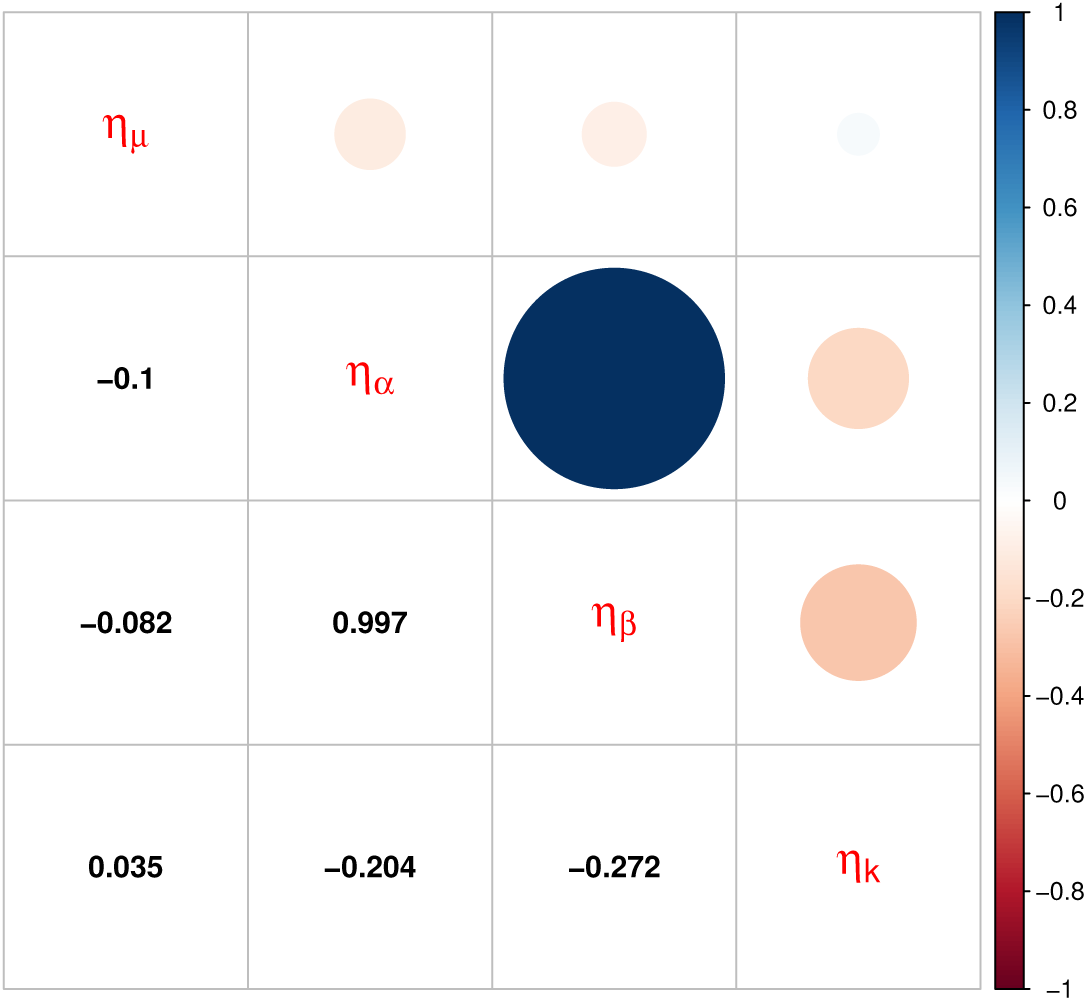
Correlations between random effects.

**Figure S8.**
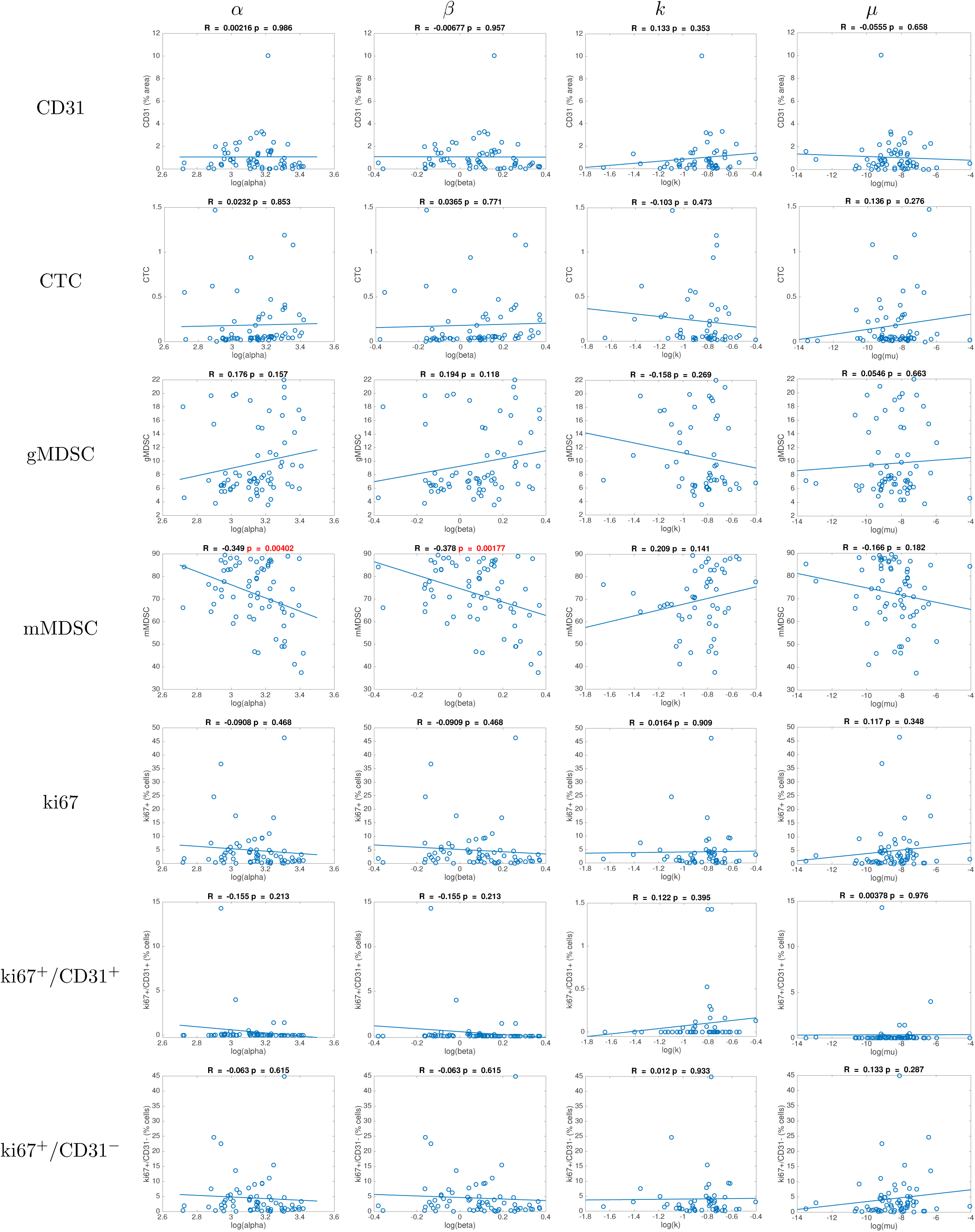
Individual parameters vs covariates.

**Figure S9.**
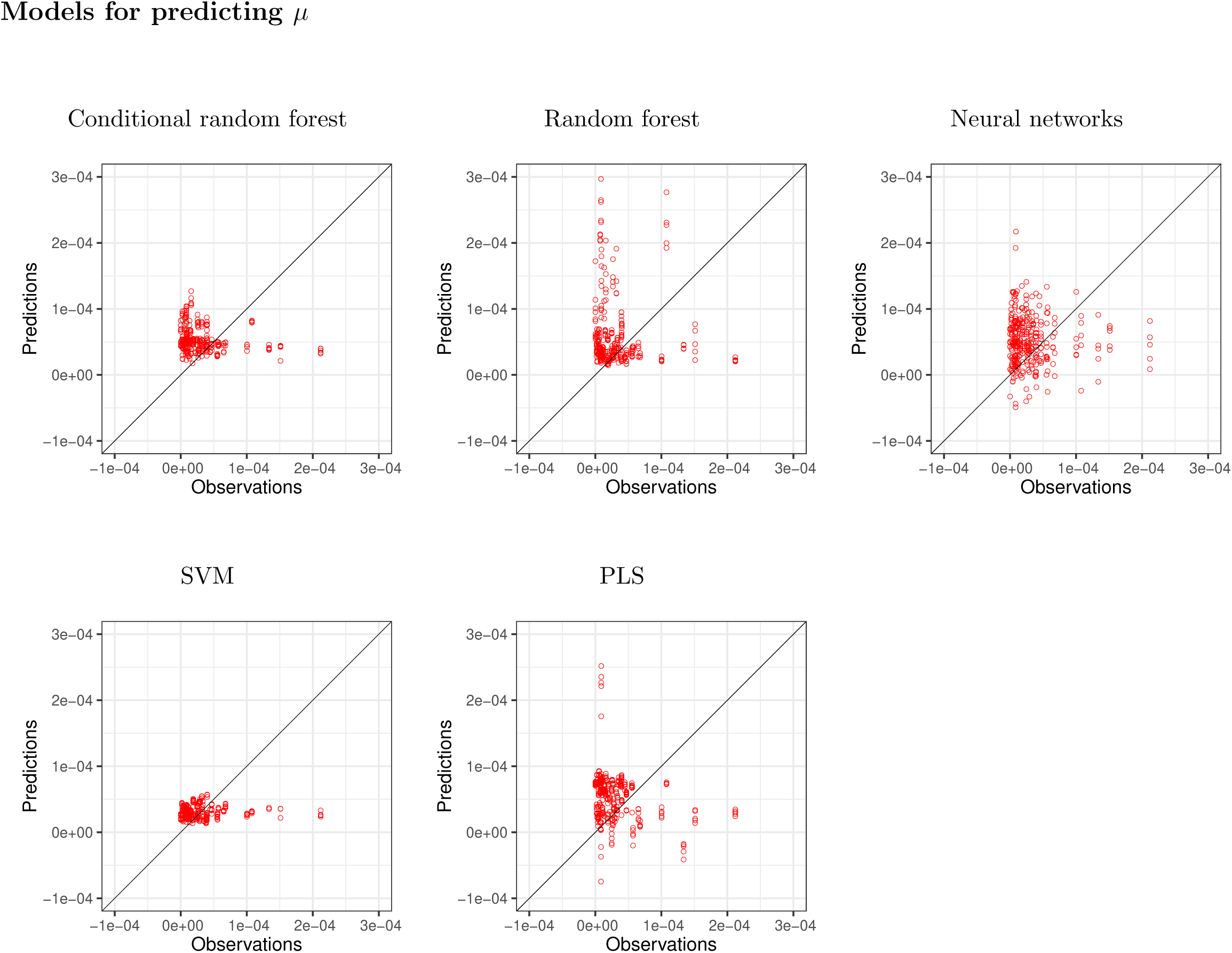
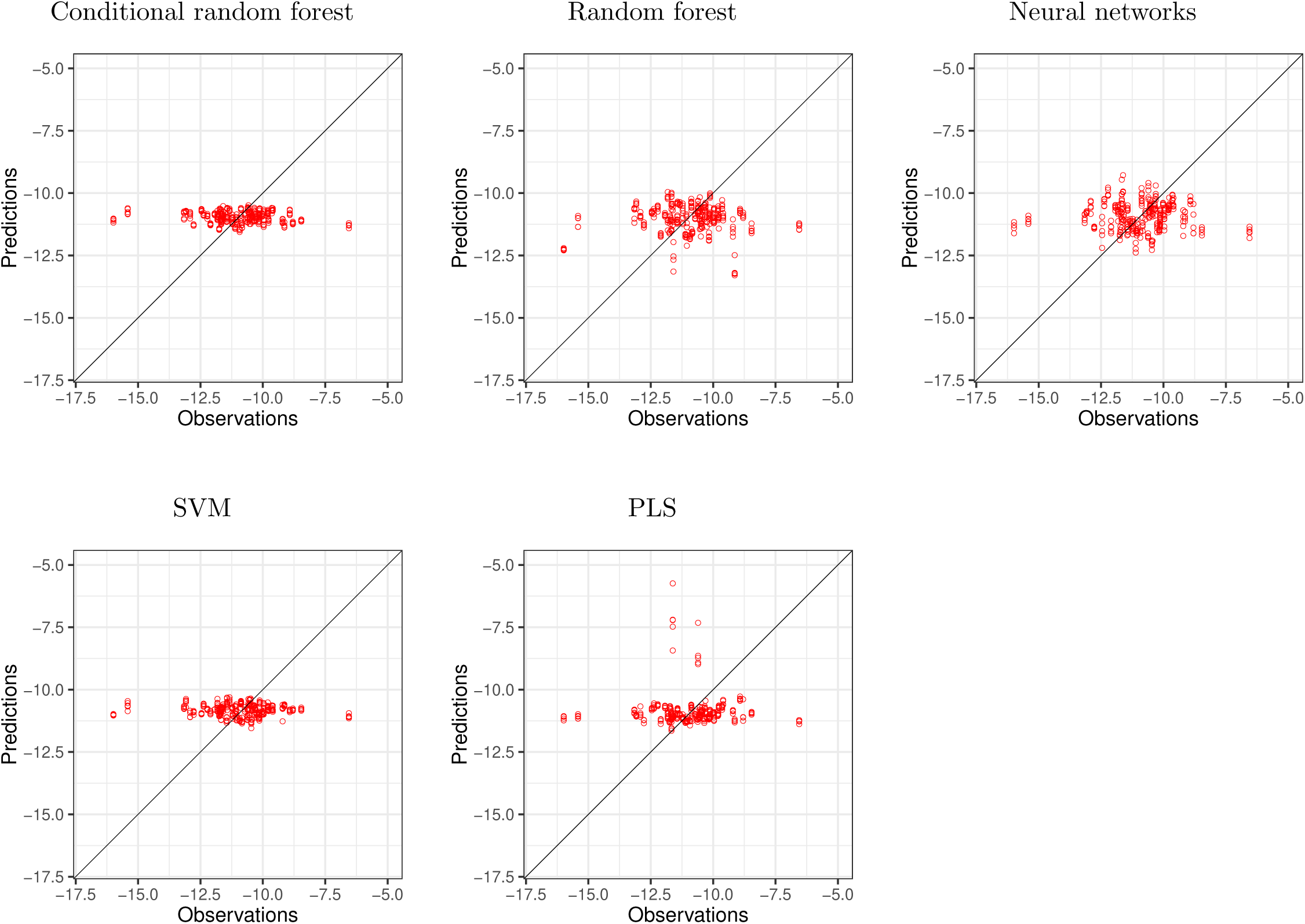
Observed vs Predicted values for the machine learning algorithms.

